# Dual reference method for high precision infrared measurement of leaf surface temperature under field conditions

**DOI:** 10.1101/2021.04.25.440729

**Authors:** Jonathan D. Muller, Eyal Rotenberg, Fyodor Tatarinov, Irina Vishnevetsky, Tamir Dingjan, Abraham Kribus, Dan Yakir

## Abstract

- Temperature is a key control over biological activities from the cellular to the ecosystem scales. However, direct, high precision measurements of surface temperature of small objects such as leaves under field conditions with large variations in ambient conditions remain rare. Contact methods such as thermocouples are prone to large errors. The use of non-contact remote sensing methods such as thermal infrared measurements provides an ideal solution, but their accuracy has been low (in the order of ~2 °C) due to necessity for corrections for material emissivity and fluctuations in background radiation (*L_bg_*).
- A novel ‘dual-reference’ method was developed to increase the accuracy of infrared needle-leaf surface temperature measurements in the field. It accounts for variations in *L_bg_* and corrects for the systematic camera offset using two reference plates.
- We accurately captured surface temperature and leaf-to-air temperature differences of needle-leaves in a forest ecosystem with large diurnal and seasonal temperature fluctuations with an uncertainty of ±0.23 and ±0.25 °C, respectively.
- Routine high precision leaf temperature measurements even under harsh field conditions, such as demonstrated here, opens the way for investigating a wide range of leaf-scale processes and its dynamics.

## 1 Introduction

Leaf pigments are optimised for the absorption of sunlight for photochemistry (Loomis, 1965). However, the heating associated with this absorption of solar energy poses a conundrum for plants: high temperatures can negatively affect a variety of bio-physical and biochemical processes (Still et al., 2019; Baldocchi and Penuelas, 2019), such as damage to photosystems I & II or the carbon reduction cycle (O’sullivan et al., 2017; Maseyk, 2006; Long et al., 1994; Werner et al., 2002). Therefore, plant canopies in high radiation and high temperature environments are challenged with the need for leaf cooling (Gates et al., 1968; Drake et al., 2018) to remain below their physiological heat limit, usually between 20–35 °C for C_3_ plants (Chaves et al., 2016; Doughty and Goulden, 2008).

Leaf cooling can partially be achieved through transpiration, leading to evaporative cooling. However, this mechanism could lead to an increased water loss due to open stomata, especially when combined with a rise in vapour pressure deficit (VPD) due to drying air or to an increase in leaf temperature (Smith et al., 2019; Richardson et al., 2020). To reduce this water loss, plants can close their stomata (Yong et al., 1997; Urban et al., 2017; Prashar and Jones, 2016), but risk a reduction in evaporative cooling. Alternatively, non-evaporative cooling relies on convection, which is controlled by wind speed and leaf structures that help adjust leaf temperature to its surrounding air (Leigh et al., 2012, 2017; Loomis, 1965). Therefore, leaf-to-air temperature difference (Δ*T_leaf−air_*) has often been used as a stress indicator in agricultural crops (Kim et al., 2018; Maimaitijiang et al., 2020; Zhang et al., 2019; Song et al., 2017; Jones et al., 2009; Long et al., 2006; Fuchs, 1990). Thus, efficient leaf cooling relies on a set of complex mechanisms to minimize Δ*T_leaf−air_*. In colder climates or seasons, leaf temperature plays different but equally important roles. For instance, leaf unfolding or frost damage (Bigler and Bugmann, 2018) could be reduced through adequate leaf temperature control. Clearly, an assessment of many physiological processes requires accurate leaf surface temperature and Δ*T_leaf−air_* measurements.

Direct and continuous measurements of leaf surface temperature remain rare in natural environments such as forests (Still et al., 2019; Aubrecht et al., 2016) and have relied on either the usage of thin-wire thermocouples or thermal infrared imagery. The usage of fine-wire thermocouples has been widespread for decades, but present multiple challenges: (a) Attaching them to leaves requires a huge effort and problems with the attachment ensue due to leaf motion, which restricts it to small sample sizes and short measurement periods (Aubrecht et al., 2016); (b) a holding structure that applies gentle pressure or some sort of glue is required, both of which can introduce errors due to their thermal conductivity and changes in surface properties (e.g. surface emissivity); (c) a large part of thermocouple junction is exposed, leading to radiative and convective heat exchange with the environment and resulting in systematic errors up to several degrees, even for tiny 0.1mm diameter junctions (Tarnopolsky and Seginer, 1999; Pieters and Schurer, 1973). Due to these difficulties, many recent studies have chosen to use infrared thermography for leaf temperature measurements (Still et al., 2019, 2021).

Infrared thermographers (IR cameras and sensors) are considered to have a low accuracy with an uncertainty of several degrees in spite of their high precision for several reasons. Firstly, systematic camera error due to instrument drift is one cause of error when cameras aren’t calibrated yearly, e.g. due to cost, as opposed to manufacturer recommendations (e.g., FLIR, 2011). For this purpose, many studies have employed a reference plate with a high emissivity to correct readings (Aubrecht et al., 2016). Secondly, the accuracy of IR temperature measurements depends on leaf/object emissivity (*∊_obj_*), reflected background thermal radiation originating from the environment (*L_bg_*), the thermal radiation emitted by the air column between leaves and the infrared camera and its transmittance. When these parameters are available, leaf/object surface temperature can be accurately calculated, either directly to the camera at the time of measurement (FLIR, 2011) or in post-processing. Inaccuracies in these parameters can lead to large measurement errors, such as changes in leaf emissivity during leaf emergence that can produce errors of up to 3 °C (Richardson et al., 2020). Also, accurately measuring *L_bg_* is challenging. Therefore, *L_bg_* has often been ignored in ecological applications, for instance due to the high emissivity of natural materials (Aubrecht et al., 2016) since the contribution of *L_bg_* becomes smaller than 5 % for *∊_obj_* > 0.95. This simplification can however lead to substantial errors in resulting temperature values (Kim et al., 2016). Direct measurements could be achieved with a hemispherical longwave sensor near the measured surface operating in the exact same spectral range as the infrared camera. However, most sensors of this type have a different spectral range than infrared cameras. Alternatively, some studies developed empirical correction equations (Kim et al., 2018) or simply used air temperature and/or relative humidity as a substitute to calculate *L_bg_* (Birami et al., 2018; Minkina and Dudzik, 2009). This technique allows for good estimates under a clear sky (Rosa and Stanhill, 2014). However, to our knowledge, the effect of these approximations has not yet been fully assessed under field conditions within the tree canopy layer. Finally, most work presently done using non-contact thermal infrared imagery in forests focused on the canopy scale or regions of the canopy (Still et al., 2019; Kim et al., 2018; Aubrecht et al., 2016; Leuzinger and Körner, 2007; Leuzinger et al., 2010), but not on the more fine-grained leaf scale.

We present a method that addresses the challenges in accurate measurements of leaf temperature (*T_leaf_*) and leaf-to-air temperature differences (Δ*T_leaf−air_*) using thermal infrared imaging in the changing environment of field conditions. This method can be applied to leaves as tiny as conifer needles in mature trees. We used an infrared camera and two reference plates to continuously and accurately measure (a) *L_bg_* and (b) *T_leaf_* and Δ*T_leaf−air_*. To demonstrate the use of our proposed method, we employed our setup in a natural forest environment.

### 1.1 Theory

Infrared thermographers measure the thermal longwave radiation sensed in their specific range (e.g., 7.5–13 μm for FLIR A320), and they are factory-calibrated to output the temperature equivalent of the full thermal range (4–100 μm; Ross, 2012). A chain of reflections and absorptions of thermal radiant fluxes (*L*; W m^−2^) affects the signal recorded by the sensor and requires consideration (Aubrecht et al., 2016; Incropera et al., 2006). The total radiant flux (*L_camera_*) received by the camera sensor is a combination of (a) the radiation emitted by the measured leaf/object *L_obj_* with emissivity *∊obj* extracted from the relevant pixels of infrared images, (b) the surface hemispherical background radiation reflected from the object *L_bg_* with reflectance 1 − *∊_obj_*, with both of these components attenuated by the transmittance *τ* of the air column between the object and the camera, and (c) the thermal radiation emitted by the air column between the object and the camera *L_air_* emitting with 1 − *τ*. These effects are integrated in Eq. 1 and illustrated in Fig. 1:

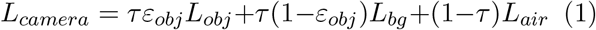

**Fig. 1.**
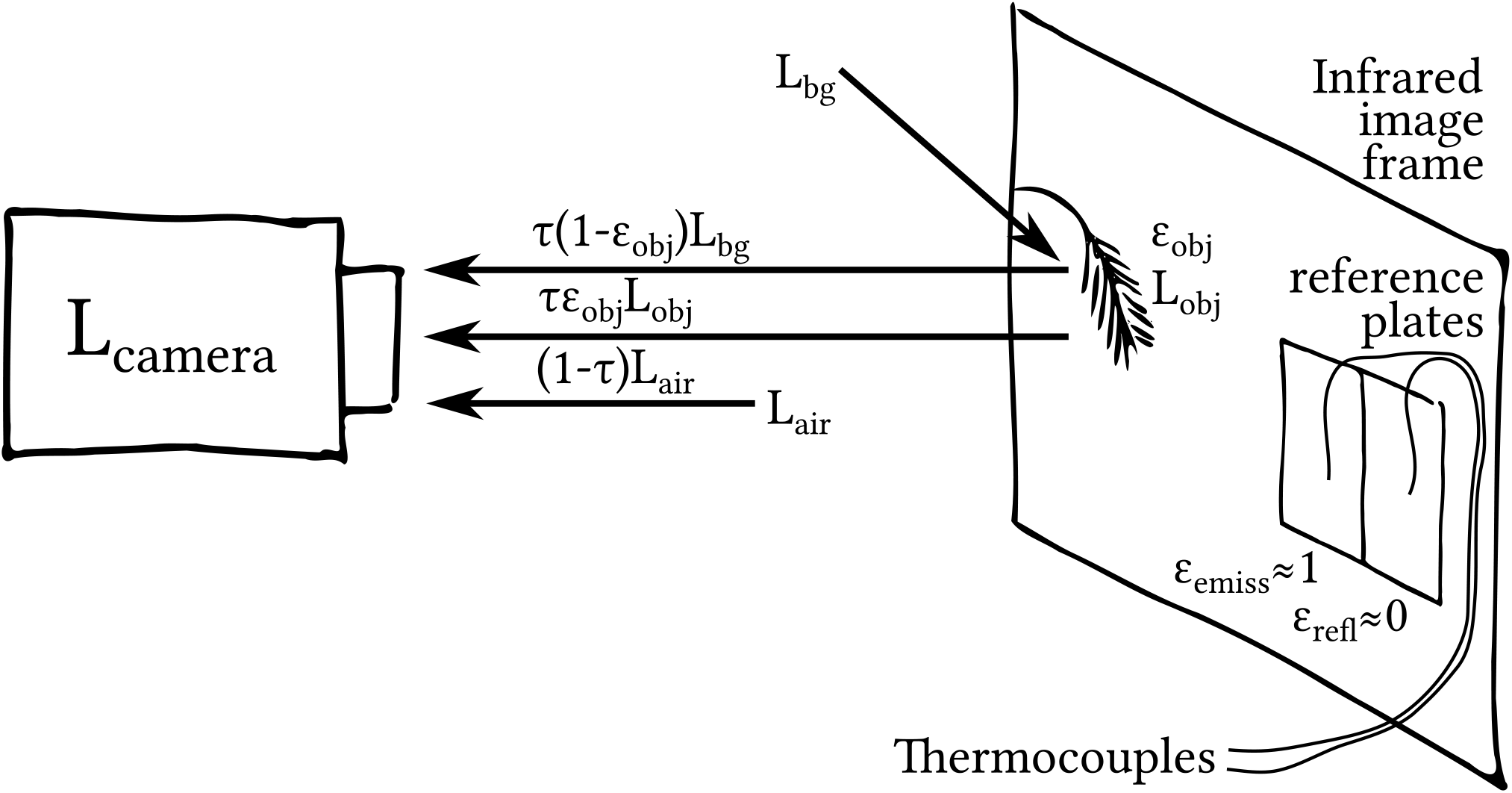
Illustration of the total radiation *L_camera_* and relevant terms received by an infrared thermographer as summarized in Eq. 1. The diagram also indicates the proposed reference plates and their built-in thermocouples (with low (*∊_refl_*) and high (*∊_emiss_*) emissivity) in the camera’s field of view that can be used to resolve the systematic camera offset and background thermal radiation (*L_bg_*), required to solve Eq. 1 for the object temperature, as demonstrated in this study

Note that in using Eq. 1, we assume a Lambertian object surface (i.e. a diffuse emission and reflection of *L_bg_* on it; Kribus et al., 2003), and no scattering due to water droplets in the air (e.g. during fog). To obtain the correct object (leaf) temperature (*T_obj_*), Eq. 1 is used together with the IR camera reading (*L_camera_*) to solve for *L_obj_*, which is then converted to temperature using the Stefan-Boltzmann law, and its constant, *σ* (Eq. 2). Note that *L_obj_* is the blackbody equivalent emitted radiation of the leaf (i.e. *∊_obj_* = 1), since *∊_obj_* is already applied in Eq. 1.

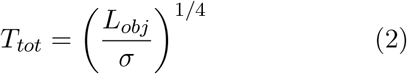

Typically, infrared cameras calculate the object temperature *T_obj_* internally using operator-provided fixed values of *∊_obj_*, *L_bg_*, *L_air_* and *τ* (FLIR, 2011). However, these variables vary significantly across time and surfaces, especially in field conditions. Accurate temperature estimates must rely in this case on post-processing of the preliminary measurements of the apparent temperature *T_ap_* (i.e., with *τ* and *∊* set to 1 in camera). To improve the uncertainty of the temperature estimate, independent information on these variables can be obtained and applied as summarized below in our ‘dual-reference’ method taking both the systematic camera offset and *L_bg_* into account.

#### Correction for the systematic camera offset

Thermal sensors suffer from systematic errors due to their sensitivity to environmental conditions (Kim et al., 2018). Kim et al. (2018) recommend using a reference plate with a high emissivity (*∊emiss* ≈ 1) and its independently measured temperature for calibration. This routine provides a corrected apparent temperature (*T_ap,cor_*) that can substitute for *T_ap_* in Eq. 1. This routine adjusts the thermal infrared readings to the reference temperature sensor used to measure this emissive plate. Note that if the final application includes a comparison to air temperature (as done here), cross-calibration of the temperature sensors of the calibration plate and the air is required to maintain high accuracy.

#### Air column variables *L_air_* & *τ*

The full equation used by infrared thermographers (Incropera et al., 2006; FLIR, 2011; Aubrecht et al., 2016) takes the air column between the camera and the object into account. *Lair* and *τ* depend on factors such as air temperature, humidity and aerosol content (Gates, 2012). At a distance of <10 m between the camera and the object, the error is −0.5 °C (Faye et al., 2016). Therefore, it is reasonable to assume *τ* ≈ 1 at a shorter distance (Aubrecht et al., 2016; Usamentiaga et al., 2014), and the right-hand term in Eq. 1 can be assumed to be practically zero.

#### Emissivity *∊*

The emissivity of the leaves and other surfaces involved in the measurements is often estimated from literature values, which can involve substantial uncertainty with a significant impact on temperature estimates (Richardson et al., 2020). Direct and high precision measurements of surface emissivity can be achieved in the lab, as we recently demonstrated using a home-made high precision system (Vishnevetsky et al., 2019), or for large enough samples in the field using the two-lid box method (Rubio et al., 2003).

#### Background radiation *L_bg_*

*L_bg_* is often ignored in infrared temperature measurements due to difficulties in measuring it near the leaves. However, as demonstrated here, it is probably a key factor in obtaining accurate temperature. Here we propose that this difficulty can be overcome by using a second calibration plate with a high reflectance (*∊_refl_* ≈ ≈ 0) in the camera field of view (Fig. 1). Estimates can be achieved by solving Eq. 1 for *L_bg_* using the camera reading of the thermal radiation originating from the reflective calibration plate, its known emissivity and its independently measured temperature.

## 2 Materials & Methods

### 2.1 Measurement Procedure

For the entire correction procedure of infrared temperature measurements, the temperature of both reference plates needs to be measured accurately and independently, e.g. using calibrated thermocouples or thermistors, thus providing *T_emiss_* and *T_refl_*, respectively. For the continued discussion of the methods, the term ‘thermocouple’ is used and we assume *τ* ≈ 1. The following steps show our method’s calculation procedure of leaf temperature (Fig. 2). Note that before any calculations can be done, the relevant pixels containing each object (leaves, reference plates) need to be identified and their apparent temperature extracted from infrared images.

**Fig. 2.**
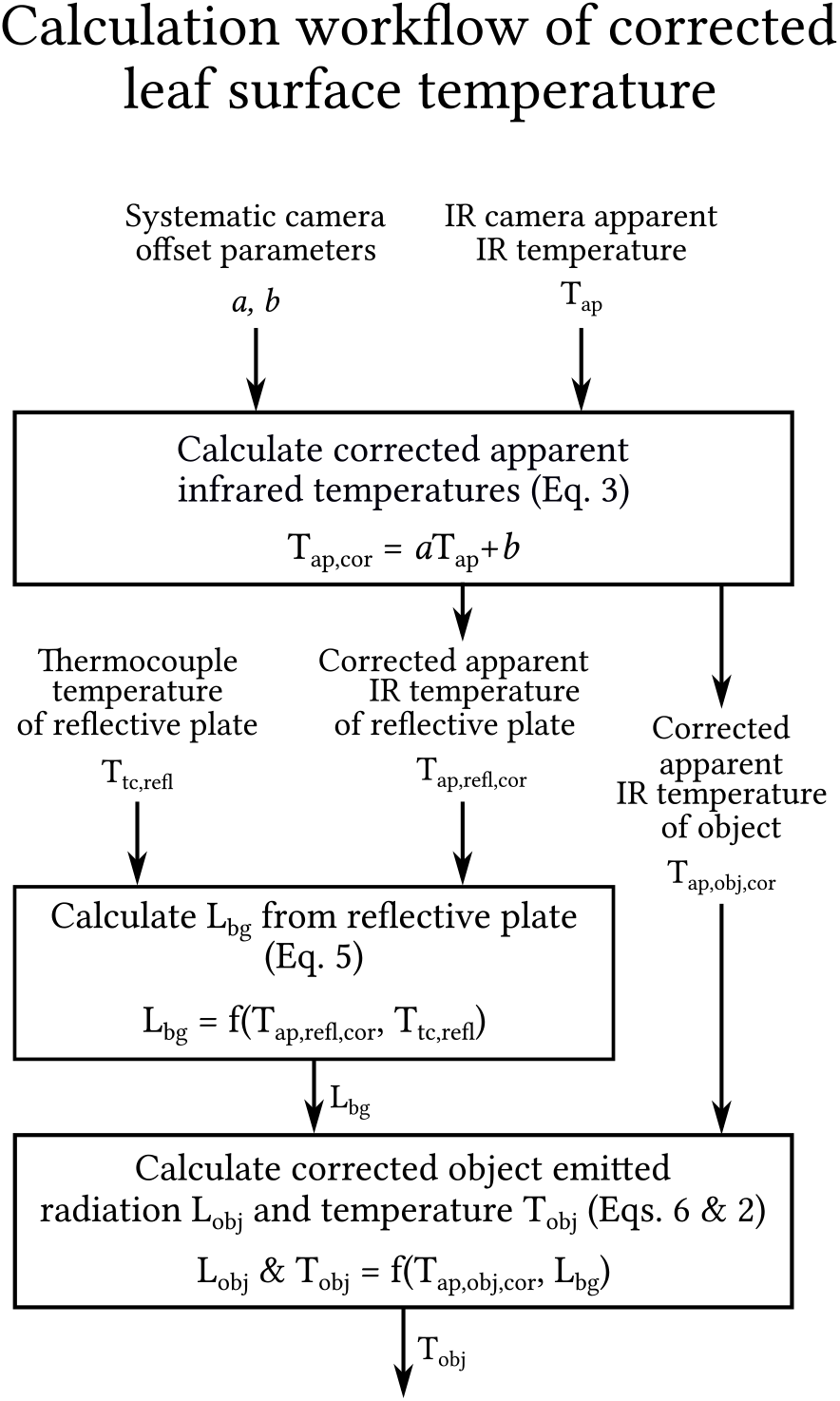
Flowchart of the calculation procedure of infrared measurements of leaf surface temperature including the measurement and correction for the effect of *L_bg_* using a reflective reference plate

#### In-situ calibration for systematic camera offset

Due to the small drift and frequent calibration, we note that the systematic camera offset follows a linear regression of the form of Eq. 3, where the corrected apparent temperature of any surface *T_ap,cor_* corresponds to the raw camera reading of the apparent temperature *T_ap_*, corrected for the camera offset using parameters *a* and *b*:

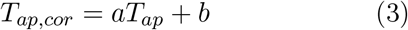

The apparent and corrected temperatures of the emissive plate are used for the calibration of the camera, and the latter depends on measurements of *L_bg_*. However, accurate measurements of *L_bg_* using a reflective plate also depend on a and b. Therefore, a simple correlation between the temperature of the emissive plate measured using a thermocouple and the infrared camera is not sufficient for calibration. Instead, a and b need to be applied to the *T_ap_* of both the reflective and emissive reference plates during the calibration process. A simple search algorithm for a and b is presented in the supporting information (Section S2) for a proof-of-concept (but this could also be done in different ways).

#### Camera-observed thermal radiation *L_camera_*

The total longwave radiation corresponding to *T_ap,cor_* measured at each pixel in the camera image, *L_camera_*, is calculated for the entire image using the Stefan-Boltzmann equation (Eq. 4), where *σ* is the Stefan-Boltzmann constant and we assign *∊_camera_* = 1.

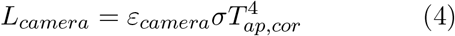

#### Thermal background radiation *L_bg_*

Next, Eq. 1 is re-arranged and solved for the background longwave radiation *L_bg_*, as shown in Eq. 5, using the thermal radiation emitted by the reflective reference plate (*L_refl_*) and received by the infrared camera (as in Eq. 4: 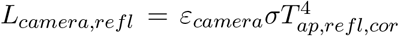). The emissivity of the reflective reference plate *∊_refl_* is known in Eq. 5, and *L_refl_* is the blackbody-equivalent radiation calculated from the independently measured reflective plate temperature *T_refl_* (using a thermocouple), i.e. 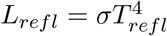.

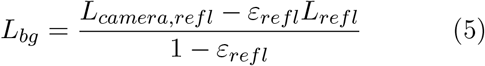

Note that in outdoors environments, it is common to assume that *L_bg_* is diffuse, but this may be unjustified, for example, when the sky’s effective temperature is much lower than ambient temperature.

#### Leaf-emitted thermal radiation *L_obj_*

Eq. 1 is then solved for the object longwave radiation *L_obj_* Eq. 6. *∊obj* is the object emissivity and 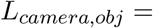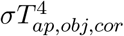 (Eq. 4).

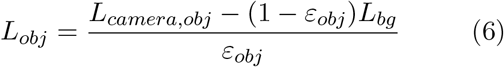

Finally, the object temperature *T_obj_* is obtained by reversing the Stefan-Boltzmann equation (Eq. 2).

### 2.2 Lab emissivity measurements

The hemispherical directional emissivity (at a right angle to the plates) of each reference plate and of natural materials present in the Yatir forest (leaves, branches, soil; Table 1) were measured in the lab at an uncertainty of <0.5 % for *∊* > 0.90 using a system developed for measurements of natural materials (Vishnevetsky et al., 2019).

**Table 1.**
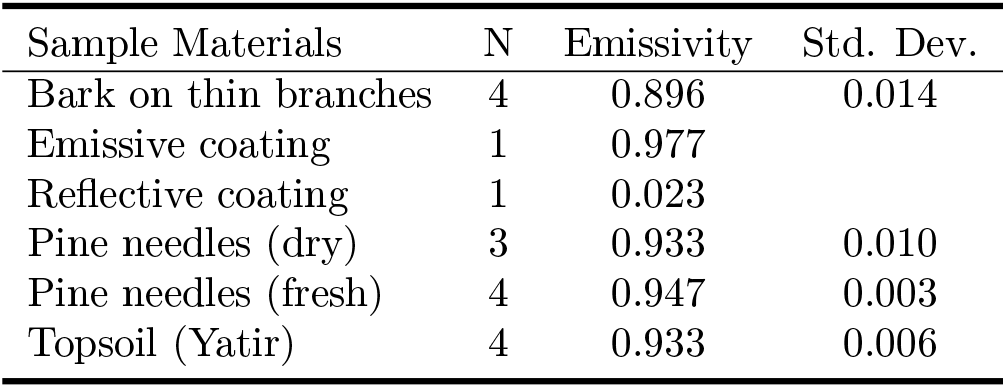
Mean hemispherical directional emissivity with number of samples (N) and standard deviation of different materials measured in the lab using the system by Vishnevetsky et al. (2019). For an emissivity *∊* > 0.90, the uncertainty is ≤0.5% of the measured *∊*, for *∊* < 0.95, the uncertainty is ≤0.4%, for *∊* ≤ 0.1, the uncertainty is ~28.6%. Note: Old fresh needles had an emissivity <0.005 higher than young fresh needles (values shown are for the mean), and differences in needle water content were insignificant.

### 2.3 Field site

All field measurements were done in the Yatir forest research site, located in a *Pinus halepensis* afforestation in the dry southern Mediterranean region, Israel (31°20′49′′N; 35°3′7′′E; altitude 650 m above sea level). Mean annual global radiation was 238 W m^−2^, while average air temperatures for January and July are 10 and 25.8 °C respectively, with a mean annual potential ET of 1600 mm, and mean annual precipitation of 285 mm (Rotenberg and Yakir, 2010; Tatarinov et al., 2016; Qubaja et al., 2019). Measurements were done between June 2018 and October 2019. The research site includes an eddy covariance flux tower operating since 2000, whose above-canopy environmental sensors (15 m agl) include air temperature (°C), RH (%), and 4 channels of radiation (W m^−2^) with up- and downwelling shortwave (0– 4000 nm; CM21, Kipp & Zonen B.V., Delft, The Netherlands) and longwave radiometers (4–50 μm; Eppley, Newport RI), both above (15 m) and below canopy (~2 m). 3D wind speed is measured at 18.7 m above ground, and ~7 m above canopy (m s^−1^; R3-100, Gill Instruments, Lymington, United Kingdom), were used for auxiliary measurements of meteorological conditions during our measurement periods.

### 2.4 Sensor Setup & Data Processing

#### IR camera deployment

A setup was developed in order to accurately measure leaf surface temperatures under field conditions. It included an infrared camera (7.5–13 μm, 320×240 px resolution; FLIR A320; FLIR Systems, Wilsonville, Oregon, United States with *τ* and *∊_camera_* were set to 1) together with emissive and a reflective reference plates. This setup was tested by deploying it in a natural environment with fluctuating environmental conditions in a pine forest in Israel for a full year of operation. To ensure that twigs with measured needles don’t move out of focus and cover entire pixels (pixels ~0.4 mm wide compared to pine needles diameters of ~0.8–1 mm), an arm extending <65 cm from the infrared camera using a 15° lens held a few twigs and the reference plates at a fixed distance from the camera (Fig. 3). This also minimised wind motion and simplified leaf detection during infrared image analysis.

**Fig. 3.**
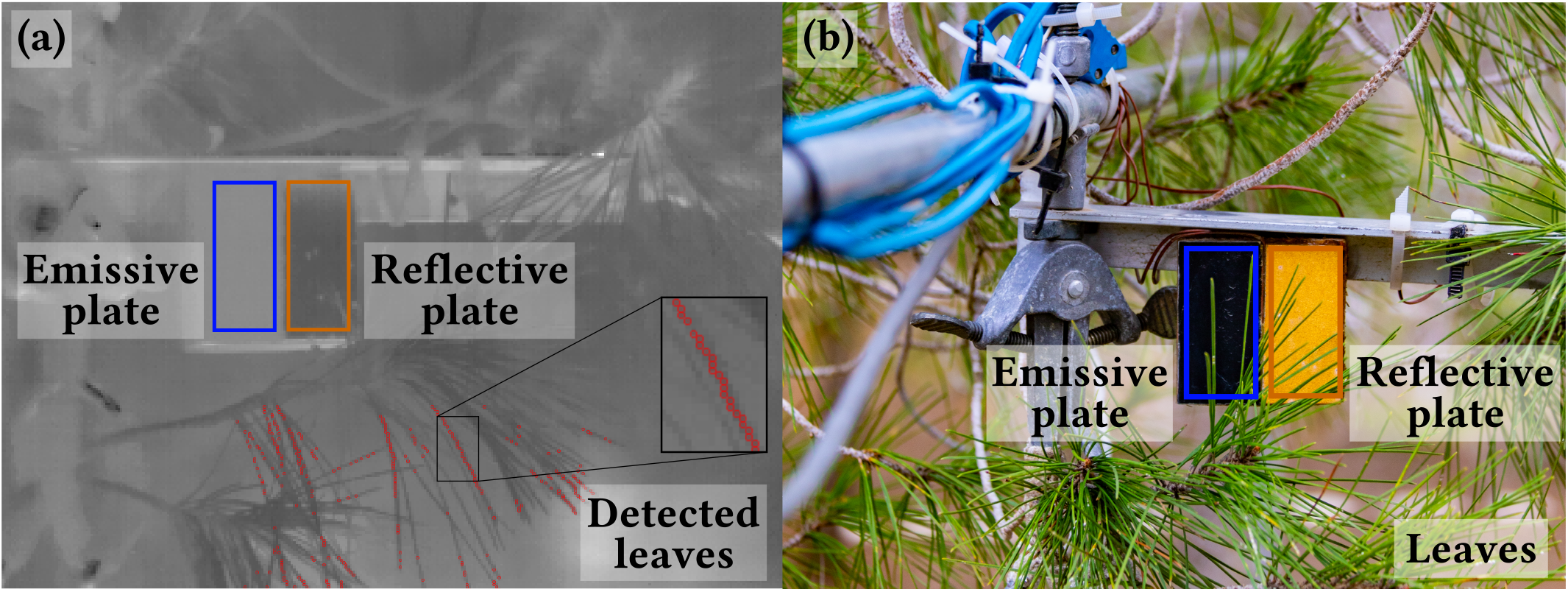
(a) Example infrared image, where red points (cf. zoomed-in square) show examples of pixels identified as leaves by our automatic script (Muller and Dingjan, 2020) and (b) setup of the infrared camera’s picture frame. Polygons show the reference plates used for camera offset correction through a known amount of emitted thermal radiation (emissive plate, blue) and background thermal radiation reflected into the IR camera (reflective plate, orange).

#### Reference plates

The two coated aluminium reference plates (plate size: 5 2.5 cm 3 mm; material chosen for its high thermal conductivity) each contained a thin thermocouple at its centre, at 0.5 mm distance from the surface, in a hole filled with thermally conductive grease drilled sideways into the plates to prevent disturbing the coated surface.

The plates were insulated by a piece of wood 2 cm thick on the back to prevent undesired heating from the environment. The emissive and reflective reference plates employed a *Metal Velvet* ™ coating (Acktar Ltd., Kiryat-Gat, Israel) and a coarse *Infragold* ^®^ coating (Labsphere Inc., North Sutton NH, USA), respectively. The coarse coating of the reflective plate produced diffuse reflection of incident background radiation, such that the image of the plate is uniform for all angles of the plate surface relative to the camera.

The reflective plate was installed vertically in order to capture *L_bg_* hemispherically, originating from above and below. Note that small differences in temperature due to fluctuations of the surface temperature of the reflective plate and the embedded thermocouple could introduce noise to *L_bg_* when the emissivity of the surface is not sufficiently low, which led to large errors when we tested it using an uncoated aluminium plate in a test (*∊* ≈ 0.3). The purpose of the embedded thermocouple is to account for the small proportion of emitted thermal radiation from the reflective plate observed by the thermal camera, as its emissivity is not exactly zero.

#### Data extraction from IR images

A matrix of raw infrared temperatures were extracted from the FLIR infrared images using a Python script developed based on an R script by Tattersal (2019). Regions of interest were scanned in horizontal lines to identify needle leaves and reference plates. A peak detection algorithm contained in SciPy (Virtanen et al., 2020) was used on each line to create a mask of valid pixels of each category: (a) To identify needles, a negative peak prominence (temperature minima) of a height of ⅓ of the total range of values in each line and a maximum width of 5 px yielded the desired leaf detection; (b) to identify reference plates, the junction of values between the ‘warm’ emissive and the ‘cold’ reflective plate was identified by calculating the first discrete difference along the horizontal line of data points, areas with a fixed width of 20 px around that junction were selected and peaks in those areas were removed (e.g. for leaves in front of the plates). The median, mean and standard deviations of all temperatures were calculated for all the pixels in each category of data (i.e. leaves and reference plates), but medians were used to reduce the effect of obvious outliers. This provided a median temperature of needle-leaves on a twig, denoted as ‘leaf’ temperature. The instantaneous data of each IR image was connected to air temperature measurements at the same time of measurement (see below).

This method limits leaf identification to conditions where the images are in focus and the background is warmer than leaves (92 % of measurements, majority of failures due to loss in camera focus), which is normally the case in our conditions due to the warm ground. To improve leaf detection, the infrared camera could theoretically be pointed up- or downward to get a clear difference to the sky or soil, but the hemispherical measurements of *L_bg_* using a reflective plate would then not provide the *L_bg_* coming from above or below. In the case of narrow needle-leaves where the leaf surface direction is not well defined, the reference plates and the camera view frame should therefore be arranged vertically. For broad-leaves, they should be arranged parallel to the side of the leaf that is measured to capture the corresponding *L_bg_*. The results of the leaf detection script were compared to manual sampling of leaf temperatures (*R*^2^ = 0.99, *P* < .001; Section S3), showing that it accurately extracts needle temperatures.

#### In-situ camera offset correction

The systematic camera offset was calibrated for the entire dataset of ~28 000 measurements using the emissive plate’s night-time temperature, as recommended by Faye et al. (2016) to achieve the optimal field measurement accuracy. Due to the heat capacity of aluminium, the thermocouple inside the reference plate reacts slightly more slowly to fast surface temperature fluctuations than the IR camera, resulting from sudden sun spots appearing on the surface of the plate. These result from changing shading during daytime due to branch motion in the canopy. The IR camera and thermocouple measurements used for calibration were highly correlated (*R*2 = 0.99, *P* < .001, *RMSE* = 0.13). Outliers resulting from sensor issues related to crashes of the infrared camera’s internal software, thermocouples, dust and humidity on the reference plates, or from rapid fluctuations of solar radiation leading to > 2±°C differences between both reference plate thermocouples were removed (<1 % of data).

#### Air temperature measurements

Thermocouple measurements of both air and reference plate temperatures were logged on a CR1000 datalogger (Campbell Scientific Inc., Logan UT, USA). For precise air temperature measurements, fine calibrated T-type thermocouples were installed in a radiation shield (Model 41003; R. M. Young Company, Traverse City MI, USA) 2–3 m away from the leaves at the same height as the camera. Calibration of all thermocouples was done in stirred water of a known uniform temperature to remove differences of individual thermocouple junctions. At ~30 °C in air, the calibrated thermocouples showed the same temperature within 0.1 °C (Section S6).

A set of fine conifer needle surface temperature thermocouples (ΔLA-C, Ecomatik, Munich, Germany) and a fine air temperature thermistor (GA10K3MCD1, TE Connectivity, Schaffhausen, Switzerland) were installed on an adjacent twig for comparison to IR readings. No correlation was found between them because thermocouple junctions were too exposed to the environmental air (see Section S4).

## 3 Results

### Camera offset correction

Infrared cameras with uncooled sensors, such as the FLIR A320, are prone to sensor drift due to environmental temperature fluctuations. FLIR mentions an inaccuracy of ±2 °C for recently calibrated cameras (FLIR, 2011). In our setup, the systematic camera offset was estimated weekly using night-time field data of the emissive reference plate and a simple search algorithm for the linear equation parameters (see Section S2). Based on this approach, a relatively large camera offset from calibrated thermocouples was estimated at 2.25 ± 0.53 °C (intercept of correction equation; slope:1.04 ± 0.03 °C. Repeating this calibration weekly indicated a small weekly drift (intercept: 0.03 ± 0.62 °C, slope: 0.002 ± 0.040 °C) around the large original correction. This data shows the importance of regular corrections for the camera offset. Notably, while here we encounter a large initial offset and relatively small subsequent drifts, the extent of such correction is likely to vary across locations and ambient conditions.

### *L_bg_* measurements

*L_bg_* was estimated using the reflective plate by solving Eq. 5 as detailed in the Methods section. These measurements are critical to improve the accuracy of the temperature estim-ates. Our values of *L_bg_* using the integrated reflective plate were compared to an *L_bg_* estimation based air temperature (using the Stefan-Boltzmann equation: 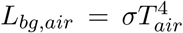) as often used when *L_bg_* measurements are not available. Under a clear sky, *L_bg,air_* consistently over-estimated *L_bg_* by up to 100 W m^−2^ (Fig. 4, *R*^2^ = 0.36, *P* < .001, *RMSE* = 27.58 W m^−2^). It improved within the canopy but still over-estimated *L_bg_* by up to ~50 W m^−2^ in most cases (Fig. 4, *y* = 0.92*x* − 14.92, *R*^2^ = 0.90, *P* < .001, *RMSE* = 13.76W m^−2^). For an over-estimation of 50 W m^−2^, resulting temperature errors can reach 0.5 °C or >1 °C for *∊* = 0.95 or *∊* = 0.90, respectively (Fig. 4), or twice that under a clear sky. Even using an empirical correlation, temperature errors could easily exceed 0.5 °C (*∊* = 0.90) due to the large range of *L_bg,air_* estimates.

**Fig. 4.**
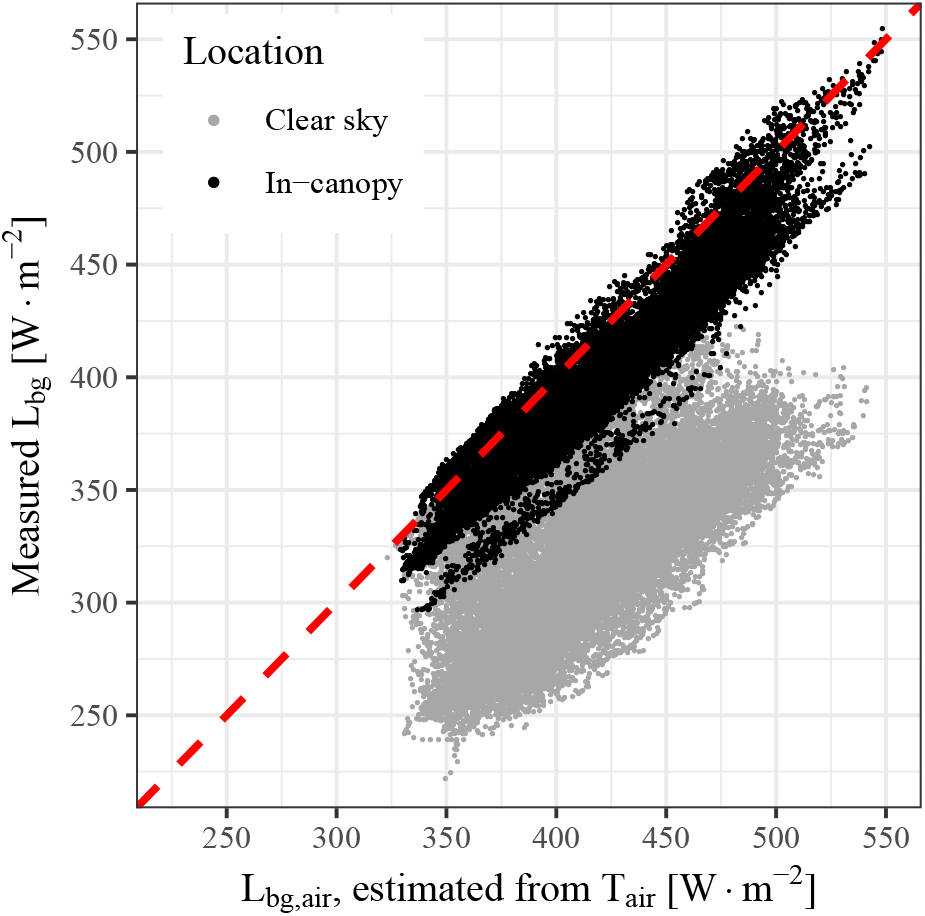
Estimates of background thermal radiation (*L_bg_*) from air temperature (*T_air_*) using a simple calculation 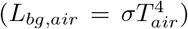, compared to measurements under a clear sky (grey) and using our reflective plate of a vertical hemisphere within the canopy (black), showing that *L_bg,air_* is consistently overestimated. Air temperature over-estimated *L_bg_* by ~50 W m^−2^ within the canopy and ~100 W m^−2^ under a clear sky

### Sensitivity to background radiation

Assessing the effects of variations in *L_bg_* on IR temperature estimates for the 300–550 W m^−2^ range naturally observed at our field site was demonstrated by the sensitivity analysis shown in Fig. 5. The temperature error could be ±1.5 °C for needle-leaves and ±3.5 °C for branches or soil, depending on the object’s surface temperature, emissivity and *L_bg_*. Note that leaf temperatures are overestimated when *L_bg_* is lower than the thermal radiation received by the camera, and underestimated otherwise. Such effects clearly limit any attempt to accurately assess Δ*T_leaf−air_* under field conditions without an accurate measurement of *L_bg_* due to its large range (e.g., during our measurement period: clear sky: 222–351 W m^−2^; vertical hemisphere in-canopy from reflective reference plate, integrated from canopy, sky and soil: 299– 552 W m^−2^).

**Fig. 5.**
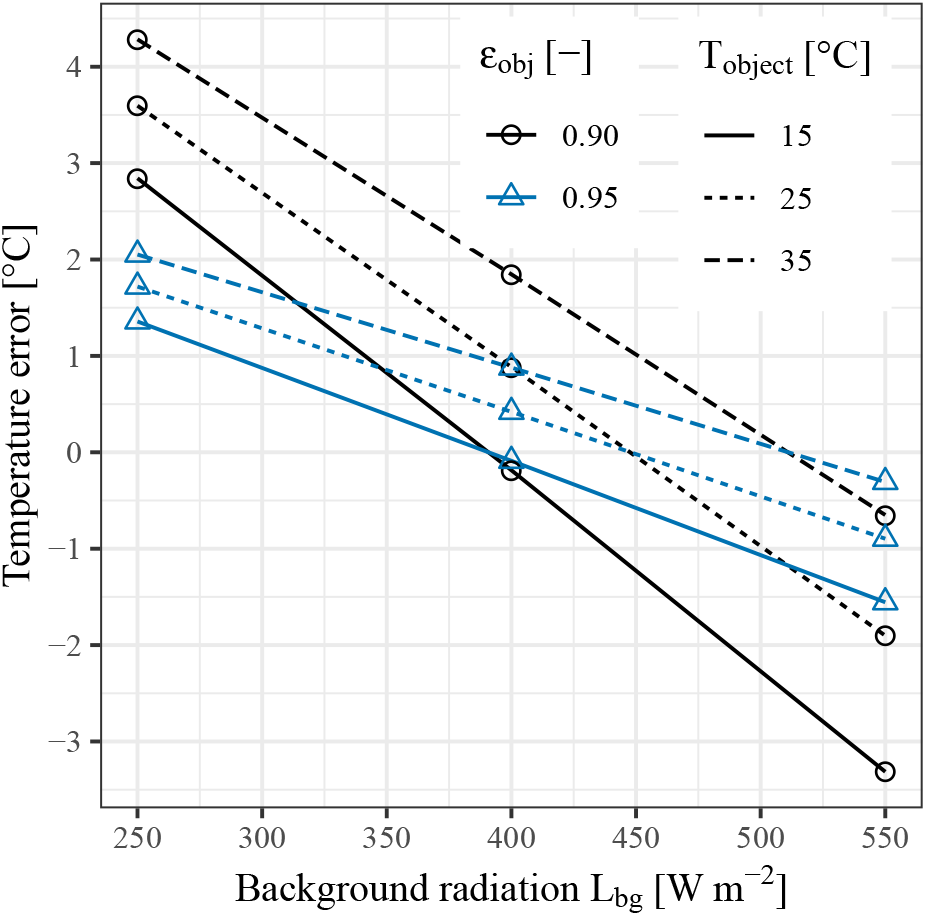
Effect of apparent object surface temperature detected by an IR camera (*T_object_*; line type), object emissivity (*∊_obj_*; colour/shape) and background thermal radiation (*L_bg_*) on infrared temperature error [°C] in field conditions, using *∊_obj_* of 0.95 (e.g. leaves) and 0.90 (e.g. branches), and *L_bg_* in field conditions ranging from 300– 550 W m^−2^, with *T_object_* ranging from 15–35 °C

### The importance of our ‘dual-reference’

While in the previous section we assessed the significance of variations in *L_bg_*, here we quantitatively assess the comparison between apparent IR temperatures readings (*T_ap_*) corrected using our method, which accounts for both the systematic camera offset and the reflected *L_bg_* (labelled as ‘Method 1’ in Fig. 6), and the previously used method using a correlation to an single emissive reference plate (‘Method 2’, according to Eq. 3), for an example of measured data (For conceptual data, see Section S7). This sensitivity analysis shows that a reference temperature (from ‘Method 1’) of 32.5 °C with *∊* = 0.90 can correspond to a 31.6–32.6 °C range for the observed *L_bg_* range when only an emissive reference plate is used (Fig. 6). For a higher emissivity of 0.95, the inaccuracy of method 2 is smaller, i.e. a range of 32.2–32.7 °C in (Fig. 6). This sensitivity analysis demonstrates the importance of taking *L_bg_* into account even for materials with a high emissivity (*∊* ≈ 0.95), such as leaves, and it shows that our method improved the measurement accuracy considerably beyond what is possible with just an emissive reference plate.

**Fig. 6.**
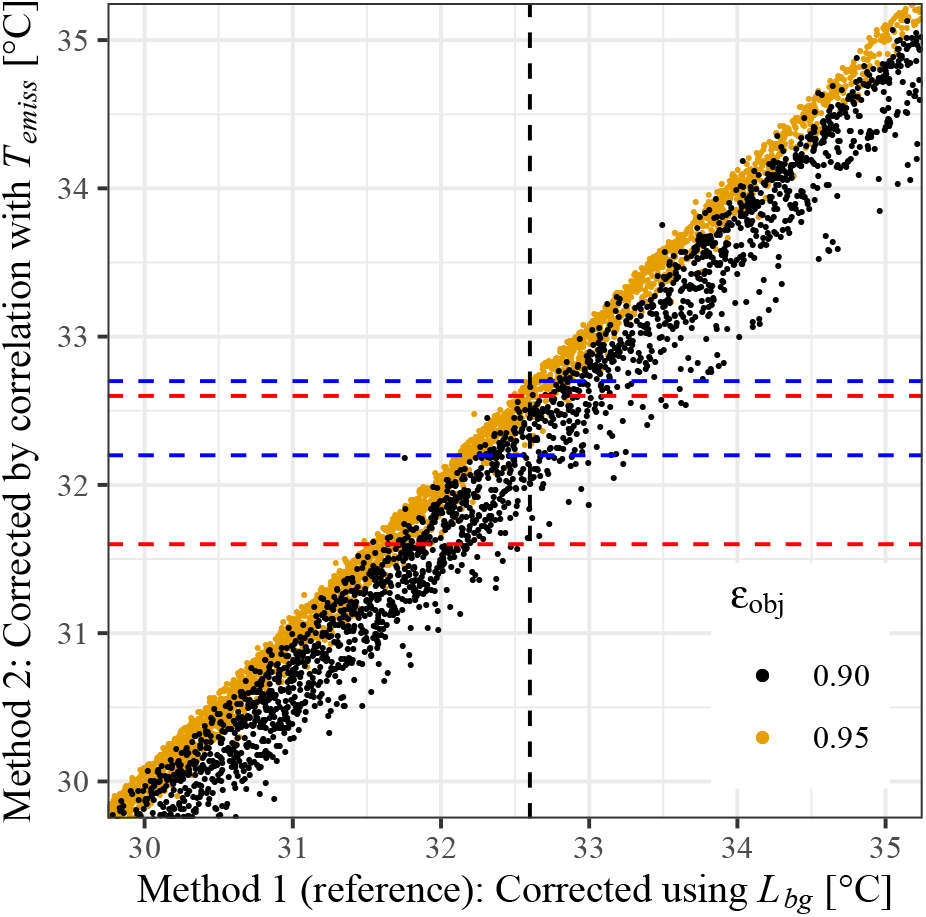
Comparison between apparent IR temperatures correction methods based on our ‘dual-reference’ method, accounting for both the reflected *L_bg_* and a systematic camera offset (x axis; ‘Method 1’, considered the reference), and using a correlation to an single emissive reference plate (y axis; ‘Method 2’; previously used), for IR measurements from 15–35 °C with an *L_bg_* of 300– 550 W m^−2^ for 2 materials of *∊* = 0.90 and 0.95 (colours). Dashed horizontal lines show the range of *T* in method 2 for a reference *T* = 32.5°C (dashed black), for *∊* = 0.90 (red) and *∊* = 0.95 (blue)

### Accuracy of leaf and leaf-to-air temperature measurements

To achieve the accuracy required to assess the leaf temperature (*T_leaf_*) and, in turn, leaf-to-air temperature differences (Δ*T_leaf−air_*) under field conditions, our ‘dual-reference’ measurement system used continuous *L_bg_* measurements and camera calibration (see Methods).

The total uncertainty of our method was calculated from the combined errors of each sensor using the log derivative method (cf. Section S1; Fritschen and Gay, 2012). Table 2 shows the resulting uncertainty of relevant variables at a leaf temperature *T_obj_* = 30 °C. The uncertainty of *T_obj_* (leaf or other object) is estimated at ±0.23 °C in our conditions (ambient temperature 10–40 °C; Fig. S1.1) and that of Δ*T_leaf−air_* to ±0.25 °C.

**Table 2.** Total resulting uncertainties of background thermal radiation (*L_bg_*), object/leaf-emitted thermal radiation (*L_obj_*), object/leaf surface temperature (*T_obj_*) and Δ*T_leaf−air_*, at object/leaf surface temperature *T_obj_* = 30 ±0.05°C, assuming and object/leaf emissivity *∊_obj_* = 0.95 ± 0.004, *∊_emiss_* = 0.95 ± 0.004, *∊_refl_* = 0.023 ± 0.007, *T_tc_* = ±0.1°C and systematic camera offset corrected by *T_ap,cor_* = 1.0221 *T_ap_ −* 1.8545

| Value | Uncertainty | Unit |
| --- | --- | --- |
| $L_{bg}$ | $\pm 3.44$ | $\text{W m}^{-2}$ |
| $L_{obj}$ | $\pm 1.32$ | $\text{W m}^{-2}$ |
| $T_{obj}$ | $\pm 0.23$ | °C |
| $\Delta T_{leaf-air}$ | $\pm 0.25$ | °C |

### Implications to field studies

The implications of our integrated ‘dual-reference’ approach to field studies of *T_leaf_* and Δ*T_leaf−air_* is demonstrated by our measurements in the study site within the canopy (in 5 m height in the middle of the tree canopy) from the summer of 2018 to that of 2019. From Nov. 2018 to June 2019, the setup was kept in the same location, but exposed to large diurnal and seasonal fluctuations, including the rainy winter to the hot and dry summer. Measurements were grouped in bins of 100 W m−2 of incoming solar radiation for the analysis (Fig. 7). Night-time temperatures were as much as 15 °C lower than daytime, and the incoming solar radiation mostly remained below 800 W m−2 due to some partial canopy shading (while above-canopy SWR reached >1000 W m^−2^). Therefore, bins above this value contained very few measurements (*n* < 20). At night, Δ*T_leaf−air_* dropped below zero, most likely due to radiative and evaporative cooling of the leaves in the absence of incoming solar radiation. The results reported in Fig. 7 demonstrate the large seasonal temperature range of about 30 °C at the field site, and the order of magnitude smaller Δ*T_leaf−air_* values of about 1.46 0.73 °C (bins >200 W m^−2^). In spite of these challenging conditions, the differences between leaf and air temperature were found to be significant in each bin in paired *t*-tests (*P* < .001; Fig. 7a) and on average, Δ*T_leaf−air_* differed significantly from zero (> 1*σ*; bins >200 W m^−2^; Fig. 7b). These results clearly show that measurements at an uncertainty of ±0.25 °C are required to capture *T_leaf_* sufficiently accurately and, in turn, the identified small but significant Δ*T_leaf−air_* values, in spite of the large variability in background conditions of this forest ecosystem.

**Fig. 7.**
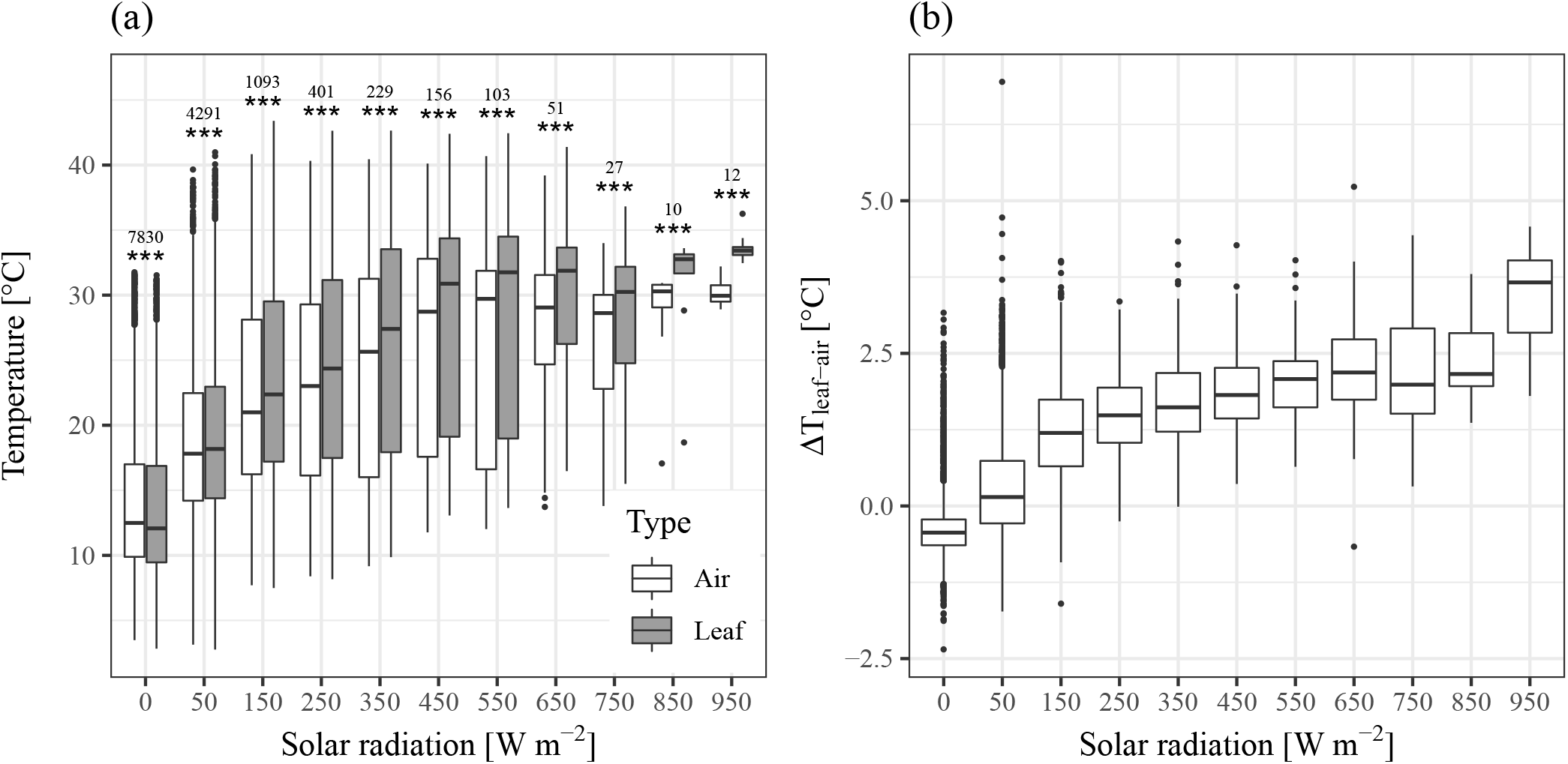
Temperatures in bins of 100 W m^−2^ of solar radiation for November 2018 – June 2019 of (a) Leaf and air temperatures with significance of differences between them and number of paired measurements, as well as (b) leaf-to-air temperature differences Δ*T_leaf−air_*. Paired *t*-tests of leaf and air temperature showed significant differences in all bins, and Δ*T_leaf−air_* is significantly larger than zero in bins >200 W m^−2^

## 4 Discussion

Accurate measurements of leaf temperature are crucial to understand many biochemical and biogeophysical processes, but remain challenging, particularly under field conditions. A number of recent studies have focussed on using thermal infrared cameras (Still et al., 2019 and references therein), but those require corrections that are difficult to make, such as for the emissivity of the measured materials (Richardson et al., 2020) and for the background radiation, which is subject to large fluctuations in natural environments (Still et al., 2019; Aubrecht et al., 2016), as well as systematic camera drift.

The results of this study identify several critical factors that limit infrared-based temperature measurements and demonstrate ways to overcome these limitations to obtain the precision required for the study of *T_leaf_* and Δ*T_leaf−air_* under field conditions. First, while direct measurement of emissivity is challenging, field methods (Rubio et al., 2003) and labscale high precision systems (Vishnevetsky et al., 2019) can be constructed and used to identify variations in *∊* values of plant materials. Second, infrared measurements are restricted by the camera accuracy without calibration, i.e. 1–2 °C (Kim et al., 2016, 2018). To reduce this error, calibration of the IR camera using an emissive reference plate is typically recommended, although that alone does not allow the error to drop below ~1 °C (Aubrecht et al., 2016). Third, significant variations in background thermal radiation (*L_bg_*) under field conditions can lead to inaccuracies of ~1.5 or ~3.5 °C in surface temperature for *∊_obj_* = 0.95 or 0.90, respectively (Fig. 5). Clearly, both the systematic camera offset and the effect of *L_bg_* must be taken into account.

A novel ‘dual-reference’ method was developed to overcome these issues in field measurements. The systematic thermal camera offset was corrected using a highly emissive reference plate with negligible effects of *L_bg_*, as previously suggested (Aubrecht et al., 2016). Our results show the importance of regular corrections for the camera offset, and while this was done weekly in the present study, a daily calibration might be advisable. Measurements of *L_bg_* were achieved using a IR camera readings of a highly reflective reference plate (7.5–13 μm). Furthermore, the correlation between these *L_bg_* measurements and a commercial full-range sensor (4–100 μm; Section S5) allows for a high level of confidence in our method to sense *L_bg_*. The temperatures of both reference plates were measured continuously using accurately calibrated thermocouples.

Measurements of non-contact leaf surface temperature has been difficult in the past because of the small differences between leaf and air temperatures (<4 °C) in the high seasonal and diurnal variability of field conditions (here: ~10–40 °C). Our system was deployed for ~2 years in rough field conditions, requiring only minimal maintenance (cleaning the camera lens and two reference plates). Using it, significant leaf-to-air temperature differences were determined across a wide range of environmental conditions. Importantly, while the magnitude of Δ*T_leaf−air_* was small, it was significantly different from zero based on the precision achieved here (Fig. 7). Since the standard deviation of Δ*T_leaf−air_* was larger than the measurement uncertainty, the level of accuracy of our system is sufficiently robust to measure *T_leaf_* and Δ*T_leaf−air_* in field conditions such as ours. Furthermore, our system provided long-term measurements at a high accuracy and spatial and temporal resolutions: with our IR camera (resolution: 320×240 px; 15° lens) and a distance to the leaves of <65 cm, pixels were ~0.4 mm wide, which allowed for measurements of conifer needles (diameter ~0.8–1 mm) or even parts thereof (Fig. 3). Our method can prove important in providing accurate field measurements for an improved understanding of leaf physiological processes and models (Muller et al., 2021, submitted).

A number of caveats or potential improvements should be noted when using thermal infrared temperature measurements: (a) While broad-leaf surface temperature measurements seem easier due to their size, their emissivity can be different on both sides. (b) If the angle of the leaf towards the camera changes, e.g. from wind, the corresponding directional emissivity has to be known to ensure accurate measurements. (c) Measurements using our proposed setup are mostly limited by the accuracy of the reference plate thermocouples and could be further improved if more precise sensors are available. (d) An aspirated radiation shield could improve air temperature measurements. (e) The distance of the camera from the leaves has to be chosen appropriately to obtain pixel size smaller than the leaf dimension (<1 mm diameter in the present study) to avoid signal contamination (e.g., from sky, soil and branches). (f) Finally, we recommend the usage of a solar radiation sensor near exposed thermocouple junctions due to the effect of sunlight on their readings.

In conclusion, we demonstrate that accurate temperature measurements (±0.23 °C for *T_leaf_* and 0.25 °C for Δ*T_leaf−air_*) can be achieved on a routine basis in field conditions even for small-dimension needle leaves, and our results demonstrate the necessity for accurate measurements of *L_bg_* near the leaves due to large fluctuations in field conditions.

## Supporting information

Complete Supplement

## Nomenclature

Symbols

σ: Stefan-Boltzman constant
*τ*: Transmissivity of the air column
ε: Emissivity
*a, b*: slope, intercept
*L*: Thermal radiant flux

Subscripts

*air*: Air column between object and IR camera
*ap*: Apparent, infrared, temperature
*bg*: Thermal background radiant flux
*camera*: Camera-received radiation
*cor*: Corrected, i.e. after calibration
*emiss*: Emissive plate (*∊_emiss_ ≈* 1)
*ir*: Infrared
*obj*: Measured object of interest
*refl*: Reflective plate (*∊_refl_ ≈* 0)

## Acknowledgements

The authors are grateful to Meir Shalev for his help with the IR camera and Revital Weic for her manual sampling of needle-leaf temperature.

This study was supported by JNF-KKL (10-10-920-19), the Israel Science Foundation (ISF#1976/17), a research grant from the *Yotam project* and the Uzi Yemin grant from the *Weizmann Institute Sustainability and Energy Research Initiative*.

## Author contributions

The study was conceived by JM, ER, IV and DY; JM developed the IR system, and carried out the measurements; JM analysed the data with help of FT and TD under the guidance of AK, ER and DY; JM, AK, ER and DY contributed to the writing.

## Code and data availability

Software scripts developed for the analysis are openly available as follows:

- Script to automatically trigger an infrared camera through a LAN interface: ‘FLIR-A320-control’ at https://doi.org/10.5281/zenodo. 4088156, reference (Muller, 2020)
- Python script to extract raw temperature data from FLIR infrared images: ‘IR-data-extraction’ at https://doi.org/10.5281/zenodo.4104314, reference (Muller and Segev, 2020)
- Script to detect pine needles and reference plates in infrared images: ‘Pine-needle-thermal-detection’ at https://doi.org/10.5281/zenodo. 4284621, reference (Muller and Dingjan, 2020)

## Notes

https://doi.org/10.5281/zenodo.4088156

https://doi.org/10.5281/zenodo.4104314

https://doi.org/10.5281/zenodo.4284621

### Competing Interest Statement

The authors have declared no competing interest.

