## Supplementary material for "Dual reference method for high precision infrared measurement of leaf surface temperature under field conditions": Complete Supplement

### Supporting Information

#### S1 Full derivation steps of the error calculations

The total measurement uncertainty was calculated from the combined errors of each sensor (Table S1.1), i.e. the thermocouples, the FLIR A320 infrared camera (FLIR, 2011), emissivity measurements (Vishnevetsky et al., 2019) and the Stefan-Boltzmann constant (Fritschen and Gay, 2012). Since the thermocouples were calibrated with the thermistor and the infrared camera was adjusted to the thermocouples, the uncertainty calculation uses the accuracy of the thermistor and the precision of the infrared camera.

**Table S1.1.** Accuracy and precision values available for all components of the system

| Sensor | Accuracy | Precision | Precision [%] |
|---|---|---|---|
| IR camera (FLIR A320) | 2.0°C or 2.0% | <0.05 °C at 30 °C | 0.17 |
| Calibrated thermocouples (T-type) | 0.2°C or 0.4% | 0.03 °C | |
| Thermistor (GA10K3MCD1) | 0.2°C | | |
| Emissivity $\varepsilon_{refl} \leq 0.1$ | | | 28.6 |
| Emissivity $\varepsilon_{refl} \geq 0.90$ | | | <0.5 |
| Emissivity $\varepsilon_{refl} \geq 0.95$ | | | <0.4 |
| Stefan-Boltzmann const. $\sigma$ | | | 0.051 |

The calculation is done using the log derivative method (Fritschen and Gay, 2012) which allows to separate the contributions of each variable by using the derivation of the logarithm of the equation, i.e. $\mathrm{d}(log\ y)\,/\mathrm{d}y = 1/y$, or consequently $\mathrm{d}(log\ y) = \mathrm{d}y/y$.

First, $T_{ap}$ was corrected for the systematic camera error by applying the linear correction Eq. 3. According to the log derivative method, a logarithm is applied Eq. S1.1 before deriving Eq. S1.2, where for a variable $X$, the precision or accuracy is $\Delta X/X$. The error of the systematic camera offset Eq. S1.3 is small since the slope factor $a \approx 1$. All temperature errors are given in Kelvin.

$$logT_{ap,cor} = log(aT_{ap} + b) \tag{S1.1}$$

$$\frac{\mathrm{d}T_{ap,cor}}{T_{ap,cor}} = \frac{\mathrm{d}(aT_{ap} + b)}{(aT_{ap} + b)} \tag{S1.2}$$

Replacing derivatives (d) with finite differences ($\Delta$), we get:

$$\frac{\Delta T_{ap,cor}}{T_{ap,cor}} = a\frac{\Delta T_{ap}}{(aT_{ap} + b)} = a\frac{\Delta T_{ap}}{T_{ap,cor}} \tag{S1.3}$$

The error of the camera-perceived longwave radiation $L_{tot}$ Eq. S1.4 was calculated from the corrected apparent temperature $T_{ap,cor}$ Eq. 4 & Eq. 3. The camera emissivity $\varepsilon_{camera} = 1$ is a setting and has no error. The effect of the atmospheric transmissivity is ignored because the camera was close to the object ($\tau = 1$).

$$\frac{\Delta L_{camera}}{L_{camera}} = \frac{\Delta\sigma}{\sigma} + 4\frac{\Delta T_{ap,cor}}{T_{ap,cor}} \tag{S1.4}$$

The error of the background thermal radiation $L_{bg}$ Eq. S1.5 was calculated based on the previous error of $L_{camera}$ Eq. S1.4 and the corrected apparent (infrared) and measured (thermocouple) temperatures of the reflective reference plate, $T_{ap,cor,refl}$ and $T_{tc,refl}$, respectively. The error of the emissivity measurement of the plate $\varepsilon_{refl}$ is also considered. Since individual errors can be both, positive or negative, we sum only absolute values to insure that the errors are combined in the most unfavourable way (Fritschen and Gay, 2012).

$$\frac{\Delta L_{bg}}{L_{bg}} = \frac{\Delta\sigma}{\sigma} + \frac{\Delta T_{ap,cor,refl} 4T^3_{ap,refl}}{T^4_{ap,cor,refl} - \varepsilon_{refl} T^4_{tc,refl}} + \left| -\frac{\Delta T_{tc,refl} \varepsilon_{refl} 4T^3_{tc,refl}}{T^4_{ap,cor,refl} - \varepsilon_{refl} T^4_{tc,refl}} \right| + \frac{\Delta\varepsilon_{refl} T^4_{tc,refl}}{T^4_{ap,cor,refl} - \varepsilon_{refl} T^4_{tc,refl}} + \left| -\frac{\Delta\varepsilon_{refl}}{1-\varepsilon_{refl}} \right| \qquad \text{(S1.5)}$$

The error of the longwave radiation emitted by the object $L_{obj}$ Eq. S1.6 was calculated from the errors of $L_{camera}$ Eq. S1.4, $L_{bg}$ Eq. S1.5 and the object's emissivity measurement $\varepsilon_{obj}$.

$$\frac{\Delta L_{obj}}{L_{obj}} = \frac{\Delta T_{ap,cor,obj} 4T^3_{ap,cor,obj}}{T^4_{ap,cor,obj} - (1-\varepsilon_{obj})L_{bg}} + \left| -\frac{\Delta L_{bg}(1-\varepsilon_{obj})}{T^4_{ap,cor,obj} - (1-\varepsilon_{obj})L_{bg}} \right| + \frac{\Delta\varepsilon_{obj} L_{bg}}{T^4_{ap,cor,obj} - (1-\varepsilon_{obj})L_{bg}} + \left| -\frac{\Delta\varepsilon_{obj}}{\varepsilon_{obj}} \right| \qquad \text{(S1.6)}$$

The error of $L_{obj}$ Eq. S1.6 was used to calculate the error of the object temperature $T_{obj}$ Eq. S1.7:

$$\frac{\Delta T_{obj}}{T_{obj}} = \frac{1}{4}\frac{\Delta L_{obj}}{L_{obj}} + \left| -\frac{1}{4}\frac{\Delta\sigma}{\sigma} \right| \qquad \text{(S1.7)}$$

Finally, the error of $\Delta_{T,leaf-air}$ could be calculated according to Eq. S1.8:

$$\frac{\Delta(\Delta_{T,leaf-air})}{(\Delta_{T,leaf-air})} = \frac{\Delta T_{leaf}}{T_{leaf} - T_{air}} + \left| -\frac{\Delta T_{air}}{T_{leaf} - T_{air}} \right| \qquad \text{(S1.8)}$$

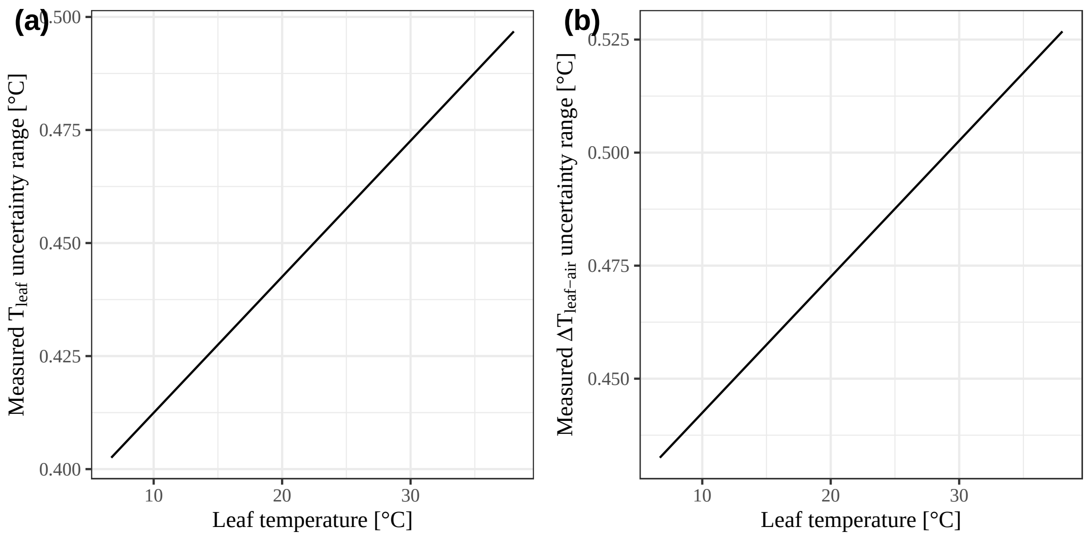


**Fig. S1.1.** Total measurement uncertainty range of leaf surface temperature ($T_{leaf}$) and leaf-to-air temperature differences ($\Delta_{T,leaf-air}$) in dependence of real leaf temperature, calculated from the combined uncertainties of all sensors used in the system.

#### S2 Calibration of systematic camera offset

When a calibration needs to be done due to sensor degradation, each physical pixel in the camera is a separate sensor and requires its own correction. A pixel noise map can be created under conditions that create good uniformity of $L_{bg}$ and temperature, as described by Aubrecht et al. (2016). The following method assumes that this sensor noise is negligible and describes a correction of the systematic camera offset.

The systematic offset, which also depends on $L_{bg}$, is assumed to follow a linear regression of the form of Eq. 3. The corrected apparent temperature of any surface $T_{ap,cor}$ corresponds to the raw camera reading of the apparent temperature $T_{ap}$, corrected for the camera offset using parameters $a$ and $b$.

A simple proof-of-concept process of calibration is described below and in the flowchart (Fig. S2.1, left). Of course, instead of the presented manual search for $a$ and $b$, a problem of numerical optimization could be devised. In this case, the objective function is the RMS difference between the measured and calculated temperatures, and $(a, b)$ is the independent parameters set.

This calibration requires a range of temperature and background radiation conditions, which can be achieved in the field using the natural variability or in the lab. Therefore, a few days of measurements are sufficient, but it will be more robust with a larger dataset. Each reference plate temperature measurement using the infrared camera is the mean of multiple pixels near the centre of the respective plate.

First, a table of parameters $a$ and $b$ (as rows and columns, respectively) in a range of $\pm 10$ with a high resolution of up to 4 decimals (for maximal precision) is created in order to test all combinations of $a$ and $b$ for the appropriate calibration values. Then, the table is filled with an initial background thermal radiation $L_{bg,initial}$ calculated for each combination of $a$ and $b$ according to Eq. 4 & Eq. 5 (by using $aT_{ap,refl} + b$ to calculate $L_{camera,refl}$).

Full calculation workflow of corrected surface temperature from infrared camera and reference plates

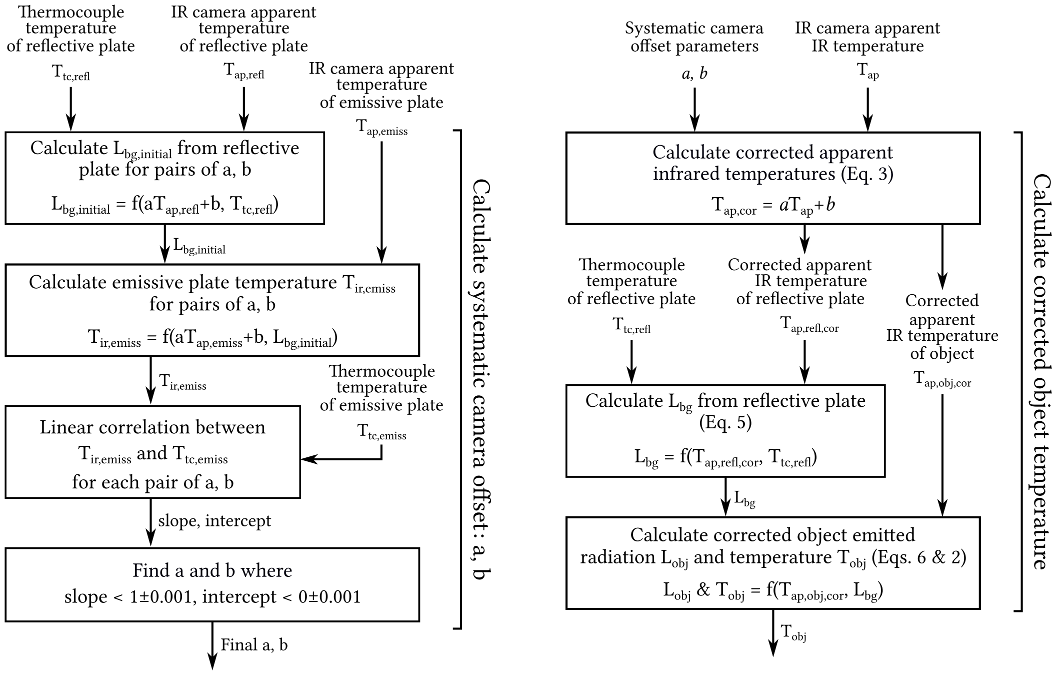


**Fig. S2.1.** Flowchart of the correction of infrared measurements of leaf surface temperature including: (left) Calculation of the systematic camera offset parameters (a, b) and (right) calculation of the $L_{bg}$ and corrected leaf surface temperature

Second, again for each combination of $a$ and $b$, a 'corrected' temperature of the emissive plate $T_{ir,emiss}$ is calculated according to Eq. 6 & Eq. 2 using their corresponding $L_{bg,initial}$, where $L_{camera,obj} = 1\sigma(aT_{ap,emiss} + b)^4$ (Eq. 4).

Third, the resulting $T_{ir,emiss}$ for all combinations of $a$ and $b$ are correlated to the independent thermocouple temperature measurements of the emissive plate temperature $T_{emiss}$. Those correlations each provide a slope and an intercept.

Finally, for the final camera calibration parameters $a$ and $b$, the infrared and thermocouple measurements should correspond to each other. Therefore, the values of $a$ and $b$ are extracted from the table where the slope is the nearest to 1 and the intercept to 0.

The values of parameters $a$ and $b$ are then used to calculate all corrected object surface temperatures $T_{ir,obj}$ according to Eqs. Eqs. (1), (2) and (4)–(6), using the apparent infrared temperatures $T_{ap.cor}$ resulting from Eq. 3 (Fig. S2.1, right).

### S3 Needle identification: Manual approach vs. script

The results of the automatic needle identification script (Muller and Dingjan, 2020) were evaluated by comparing them to manual sampling of needles contained in regions of interest (ROI) on infrared images. The two methods were highly correlated ($R^2 = 0.99$, $P < 001$; Fig. S3.1), which confirms the robustness of our script. Manual sampling involved selecting small polygons (ROI) on the image, of which the mean temperature of all pixels was calculated. The ROIs were often more than 5 px wide due to the resolution of the image and were selected in areas with numerous densely packed and overlapping needle-leaves in order to avoid sampling the unfocused background. However, this manual approach was not perfect, which reduced the exactitude of the needle-leaf selection. The automatic script was able to more precisely identify needle-leaves due to its ability to precisely identify them.

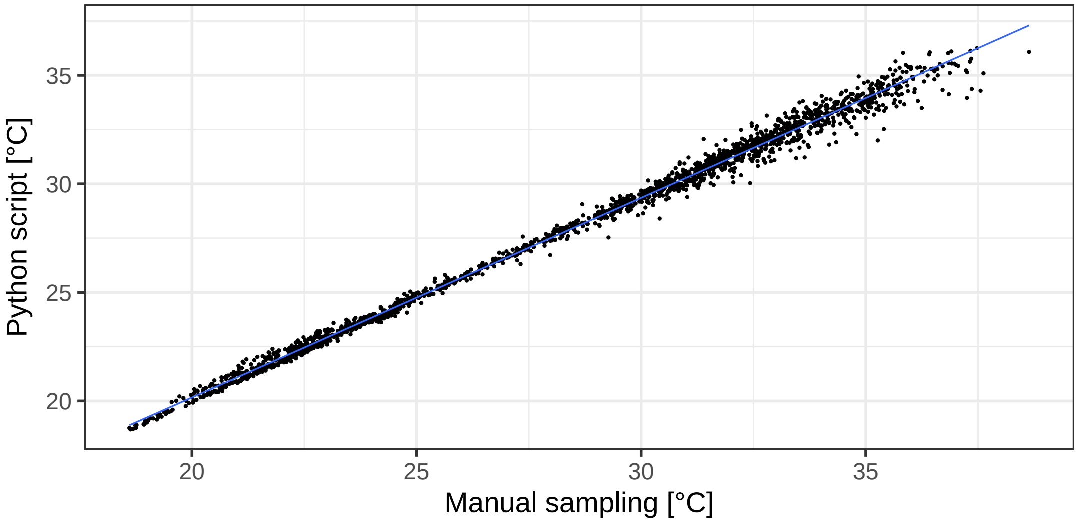


**Fig. S3.1.** Comparison between manual sampling of leaf temperature (by Revital Weic) and the automatic python script, where $y = 1.08x - 1.76$, $R^2 = 0.99$, $P < .001$. Manual sampling typically included polygons more than 5 pixels wide, which introduces small errors

### S4 Leaf thermocouples vs. infrared data

A set of fine conifer needle surface temperature thermocouples (ΔLA-C, Ecomatik, Munich, Germany) and a fine air temperature thermistor (GA10K3MCD1, TE Connectivity, Schaffhausen, Switzerland) were installed on an adjacent twig for comparison to IR readings. Air temperature measurements of the thermocouple were good (Fig. S4.1a), as shown by the correlation with the reference thermistor, both in the radiation shield ($y = 0.99x + 0.0003$, $R^2 = 0.99$, $P < .001$) and in the open air near the leaves ($y = 0.99x + 0.06$, $R^2 = 0.99$, $P < .001$). Validation of the measurements of $\Delta_{T,leaf-air}$ using our system based on an infrared camera ($\Delta T_{IR}$) with concurrent measurements using the thermocouple-based Ecomatik ΔLA-C system ($\Delta T_{tc}$) failed to show a correlation between the two methods ($\Delta T_{IR} = -2.84\Delta T_{tc} - 0.29$, $R^2 = 0.18$, $P < .001$; Fig. S4.1b). This could be explained by the difference in leaf surface measurements, where the thermocouple junctions of the Ecomatik ΔLA-C system are more exposed to the environment (Tarnopolsky and Seginer, 1999; Pieters and Schurer, 1973) compared to the infrared measurements of leaf surface temperature.

Figure S4.1 shows the correlation between the ΔLA-C conifer needle surface temperature thermocouples system (Ecomatik, Munich, Germany) with its fine air temperature thermistor (GA10K3MCD1, TE Connectivity, Schaffhausen, Switzerland) and our infrared camera readings.

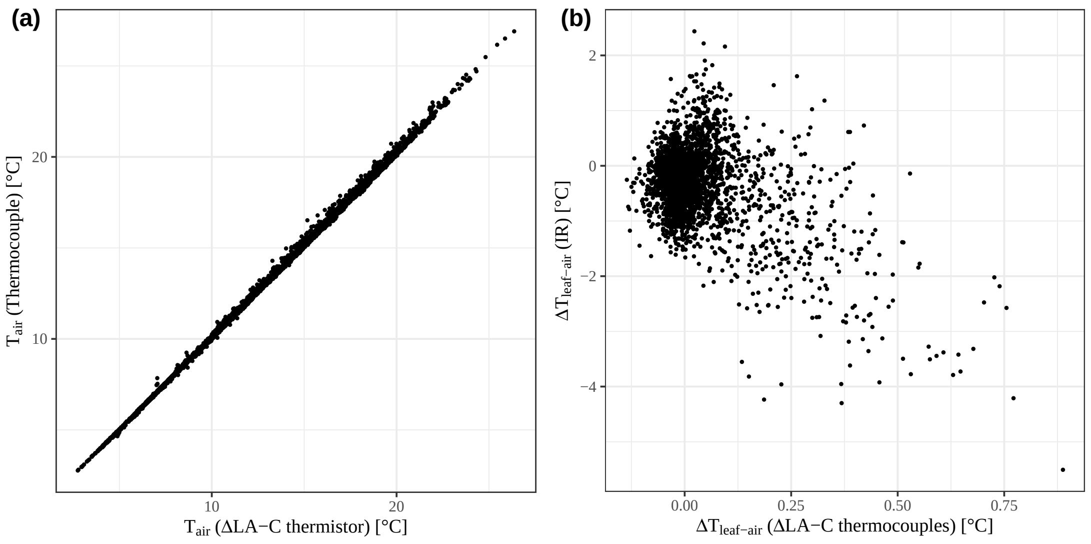


**Fig. S4.1.** Comparison between contact thermocouple-based Ecomatik ΔLA-C and infrared (IR) measurements of (a) air temperature ($T_{air,tc} = 1.02T_{air,\Delta LA\text{-}C} - 0.09$, $R^2 = 0.99$, $P < .001$) and (b) $\Delta_{T,leaf-air}$ ($R^2 = 0.18$, $P < .001$)

#### S5 Comparison of $\mathbf{L_{bg}}$ between the reflective plate and commercial sensors

$L_{bg}$ was estimated using the reflective plate by solving Eq. 5, as detailed in the Methods section (spectral range: 7.5–13 µm). Figure S5 shows a comparison to the below-canopy full-range LWR Eppley sensors 20m away from it near the eddy covariance flux tower (spectral range: 4–50 µm, calibrated to 4–100 µm) using night-time data. This comparison is not a calibration but rather a comparison to assess the consistency of the sensors due to their different spectral range. Indeed, measurements were well correlated ($R^2 = 0.98$, $P < .001$, $RMSE = 4.35$), which shows that $L_{bg}$ from IR camera measurements of the reflective plate represent the full spectral range of 4–100 µm well. The range and RMSE shown in this correlation are due to the differences in location and sensors between the two setups: The Eppley sensor is hemispherical and average measurements from a wide variety of angles (and thus area).

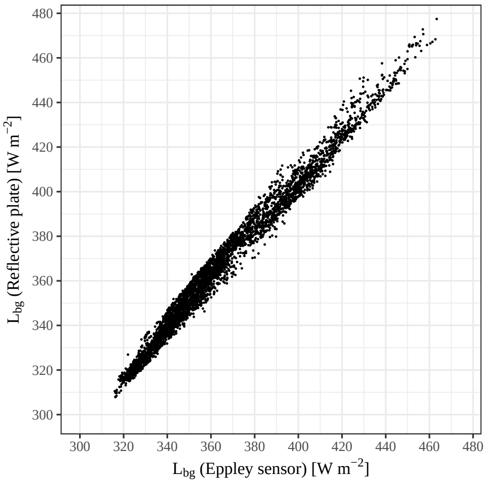


**Fig. S5.1.** Comparison between night-time $L_{bg}$ measurements using industrial Eppley sensors (4–50 µm; x axis) and the reflective reference plate using an IR camera (7.5–13 µm; y axis). Measurements are well correlated ($R^2 = 0.98$, $P < .001$, $RMSE = 4.35$). Range is attributed to different locations of both setups, but shows that the limited spectral range of the IR camera represents the full range well.

#### S6 Thermocouple calibration

The thin-wire thermocouples were calibrated in stirred ice water (to determine the offset from 0 °C). Fig. S6.1 shows the comparison between them after this calibration, under fluctuating field conditions (solar radiation, wind, etc.) while installed in a radiation shield (Model 41003; R. M. Young Company, Traverse City MI, USA). Small fast turbulent air flow fluctuations still led to minor differences between the thermocouples, but in general, their readings agreed well with each other (Fig. S6.1).

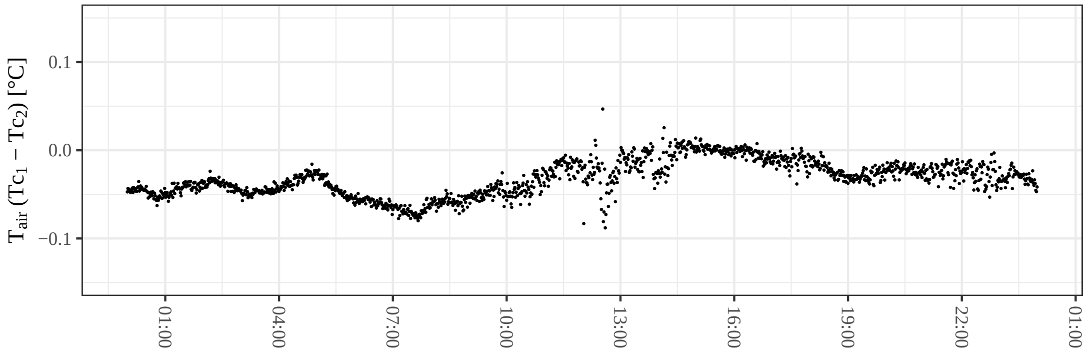


**Fig. S6.1.** Fluctuations of the difference between two thin-wire thermocouples compared to each other under field conditions in a Young radiation shield for one day with low environmental influences (e.g. turbulent wind, radiation)

### S7 Method comparison: Dual vs. Single reference plate

We quantitatively assessed including both the camera drift corrections and the effect of the reflected $L_{bg}$ using field measurements by comparing apparent IR temperatures correction methods using conceptual data ranging from 15–35 °C with an $L_{bg}$ of 300–550 W m$^{-2}$. 'Method 1' accounts for the reflected $L_{bg}$ and a systematic camera offset and 'Method 2' uses a correlation to an single emissive reference plate. A reference temperature (from 'Method 1') of 32.5 °C with $\varepsilon = 0.90$ can correspond to a 31.6–32.6 °C range when only an emissive reference plate is used (Fig. S7.1). For a higher emissivity of 0.95, the inaccuracy of method 2 is smaller, i.e. a range of 31.2–32.6 °C (Fig. S7.1).

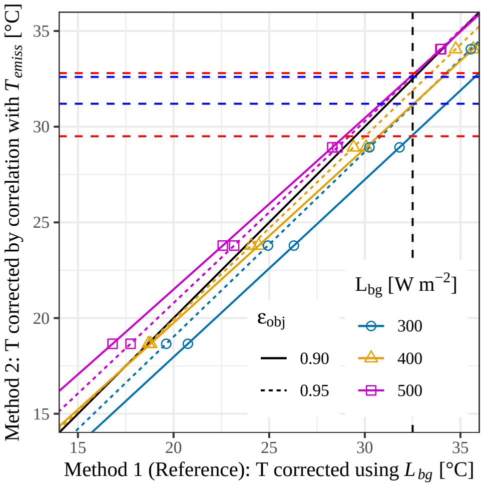


**Fig. S7.1.** Comparison between apparent IR temperatures correction methods accounting for the reflected $L_{bg}$ and a systematic camera offset (x axis; 'Method 1', considered the reference), and using a correlation to an single emissive reference plate (y axis; 'Method 2'), for conceptual data ranging from 15–35 °C with an $L_{bg}$ of 300–550 W m$^{-2}$ (colours/shapes) for 2 materials of $\varepsilon = 0.90$ and 0.95 (line types). Dashed horizontal lines show the range of T in method 2 for a reference $T = 32.5$ °C (dashed black), for $\varepsilon = 0.90$ (red) and $\varepsilon = 0.95$ (blue)
